# Information theory characteristics improve the prediction of lithium response in bipolar disorder patients using an SVM classifier

**DOI:** 10.1101/2022.04.04.486856

**Authors:** Utkarsh Tripathi, Liron Mizrahi, Martin Alda, Gregory Falkovich, Shani Stern

## Abstract

Bipolar disorder (BD) is a mood disorder with a high morbidity and death rate. Lithium (Li), a prominent mood stabilizer, is fully effective in roughly 30% of BD patients. The remaining patients respond partially or do not respond at all. Another drug used to treat BD patients is valproate (VPA). Plenty of efforts has been made to understand how these drugs affect the patients’ neurons. We have performed electrophysiological recordings in patient-derived dentate gyrus (DG) granule neurons for three groups: control individuals, BD patients who respond to Li treatment (LR), and BD patients who do not respond to Li treatment (NR). The recordings were analyzed by the statistical tools of modern information theory, which enabled us to recognize new relationships between the electrophysiological features. These added features included the entropy of several electrophysiological measurements and the mutual information between different types of electrophysiological measurements. Information theory features provided further knowledge about the distribution of the electrophysiological entities, which improved basic classification schemes. These newly added features enabled a significant improvement in our ability to distinguish the BD patients from the control individuals (an improvement from 60% accuracy to 74% accuracy) and the Li responders from the non-responders in the BD population using Support Vector Machine (SVM) classification algorithms (an improvement from 81% accuracy to 99% accuracy). These new tools showed that LR neurons are less distinguishable from control neurons after Li treatment but not after VPA treatment, whereas NR neurons become more distinguishable from control neurons after Li treatment.

## Introduction

Bipolar disorder (BD) is a severe psychiatric disorder characterized by abnormal mood episodes, typically mania and depression. During the mania state, people feel euphoric or irritable, grandiose, with increased energy and decreased need for sleep. In the depression state, they are sad and may feel empty, with associated changes in energy, cognitive functions, sleep, and appetite^1,2^.

Early diagnosis and proper long-term treatment are critical since BD illness can lead to suicide^3,4^ and typically affects education, occupational status, family connections, social interactions, and quality of life. Different pharmacological and psychosocial therapies are being used to treat this disorder. However, these have been limited in their influence on the patients^5^. The popular mood stabilizer lithium (Li) has shown a therapeutic effect in about one-third of BD patients with prophylactic treatment^6–8^ ((these patients will be abbreviated as Li Responsive (LR) patients throughout the text). The remaining patients either respond partially or do not respond at all^9,10^. We will denote these patients as Non-responsive (NR) patients throughout this study. Several studies have examined Li mechanisms of action, but these mechanisms generally remain elusive despite the efforts to understand them.

Many Li response prediction studies have used different clinical variables and features such as patients’ age, family history of BD, or the severity of symptoms^11–14^ However, the success of these prediction attempts was only partial. In 2018 Stern et al. used a naïve Bayes classifier trained on distinct electrophysiological features of LR and NR dentate gyrus (DG) granule neurons differentiated from iPSCs of BD patients for classification with a 92% success rate^15^• There have been studies to understand the underlying physiological and molecular differences between LR and NR neurons to find mechanisms for their different responses to the Li treatment. Differences in the autosomal neuronal functions of LR and NR neurons were reported in 2020 when Stern et al. showed that *KCNC1* and *KCNC2* were overexpressed in CA3 hippocampal pyramidal neurons obtained from LR BD patients (but not NR CA3 neurons). Specific potassium channel blockers were used to reduce hyperexcitability. Chronic Li therapy reduced the hyperexcitability of CA3 neurons derived from the LR BD patients. The Li treatment increased sodium currents and reduced fast potassium currents^16^.

Another study revealed that DG and CA3 pyramidal hippocampal neurons obtained from NR BD patients had an inherent physiological instability, causing rapid changes in excitability states^17^. In the most recent study, this team discovered that the activity of the Wnt/ β-catenin signaling pathway was significantly hampered in NR neurons, with a substantial decrease in *LEF1* expression. Li inhibited GSK-3 β and released catenin, which comprises a nuclear composite with *TCF/LEF1*, triggering the Wnt/ β-catenin transcription program. As a result, it was hypothesized that downregulation of *LEF1* might account for Li resistance in NR neurons^18^. Apart from this, several molecular mechanisms have been proposed to explain the cellular mechanism of Li action. Li was shown to alleviate calcium channel and mitochondrial dysfunction that were linked to various psychiatric conditions. Schlecker et al. showed overexpression of neuronal calcium sensor-1 (NSC-1) in PC12 cells and increased intracellular calcium release due to the phosphoinositide signaling pathway activation. It is well established that this family of calcium receptors in neurons regulates Ca2+ signaling^19^. Confocal microscopy was used to quantify intracellular calcium in PC12 cells. Li treatment of PC12 cells was found to inhibit the effects of NSC-1^20^. Chen et al. showed that Li raised the mitochondrial Bcl-2 expression on the outer membrane. Chronic Li and VPA therapy resulted in elevated Bcl-2 levels in the frontal cortex. Bcl-2 is a protein that acts as a neuroprotective factor^21^.

Given a high percentage of patients with low responsiveness to Li therapy, it is essential to identify and diagnose those BD patients at an early stage of the disease to avoid prolonged ineffective treatments. A few studies aim to predict the Li response of BD patients using various techniques and algorithms. A study conducted in 2002 proposed that epileptiform EEG abnormal behavior should be investigated further as a potential marker of Li resistance in BD^22^. In 2003, Washizuka and his team identified a connection between mitochondrial DNA (mtDNA) 5178 and 10398A polymorphisms and BD. A logistic regression analysis indicated that patients with the 10398A polymorphism responded considerably better to Li (p=0.03)^23^. In 2007, Li’s response was suggested to have multi-genetic etiology that relies on other clinical co-diagnoses^24^. Kafantaris and his team reported that changes in the left cingulum hippocampus (CGH) fractional anisotropy (FA) could act as a biological marker of early therapeutic efficacy. Significant differences were found when comparing variations in white matter microstructure in the CGH between LR and NR patients. LR patients’ mean CGH FA increased by 5% from the baseline at week 4 of Li treatment, while NR patients’ CGH decreased by 0.82 %^25^.

Anticonvulsant valproic acid (VPA) also has been found beneficial in some BD patients and is commonly used as an alternative to Li^26,27^. VPA was found to enhance the region of growth cones in cultured sensory neurons. After VPA chronic treatment at 0.3-0.6mM concentration, William et al. reported a decline in the rate of collapse and enlargement of growth cones in sensory neurons generated from newborn rat dorsal root ganglia^28^. According to several studies, VPA operates by engaging with the manipulation of voltage-gated sodium channels. Van den Berg et al. employed cultured hippocampal neurons to record fast spatial modulation of membrane voltage during whole-cell voltage-clamp measurements of Na+ currents. After applying 1mM VPA, a decrease in the reactivation of Na+ currents was observed. Furthermore, VPA decreased the peak Na+ conductance in a voltage-dependent manner^29^. VPA increases β-catenin transcription in the Wnt signaling pathway. Wang et al. observed that neural stem cells (NSCs) generated from embryonic Sprague-Dawley rats treated with VPA-containing media exhibited higher Wnt-3α and β-catenin than the control group. They discovered higher Wnt-3α and β-catenin expression in NSCs handled with 0.7mM VPA relative to media using RT-PCR. These findings indicated that the Wnt signaling pathway activation triggers VPA-induced neuronal differentiation^30^. In another study, VPA was shown to act downstream of GSK-3β. It upregulated LEF1 and Wnt / β-catenin gene targets, increased the complex β-catenin/TCF/LEF1 transcriptional activity, and decreased excitability in NR neurons^18^.

In this study, we use Information theory to analyze the electrophysiological recordings of BD DG granule neurons. Adding this analysis gives us new perspectives on how Li and VPA affect DG neurons of BD patients compared to the control group. Furthermore, using information theory enhances the prediction of drug response significantly. Information theory incorporates probabilistic reasoning and representation to comprehend the enriched transmission of information and processing between systems. Information theory is a handy technique even in the neuroscience field. Researchers used information theory to decode the information from neuronal populations^31,32^ previously. In 1981, it was shown using elementary mutual information theory that matching of interneurons’ contrast-response function in the compound eye of flies with the range of contrasts found in nature enables the neurons to encode contrast fluctuations most efficiently^33^. Transfer entropy, an information-theoretic quantity that quantifies linear and non-linear interactions, was previously used to systematically measure the communication between individual neurons at various time scales in cortical and hippocampal slice cultures to analyze multiplex networks of individual neurons with time scale-dependent connections^34^. Another study demonstrated the transfer entropy (TE) as a metric for efficient connectivity of electrophysiological measurements in a simple motor task-based on simulations and magnetoencephalographies (MEG)^35^. There have been other studies^36,37^ that used information theory with electroencephalography (EEG), magnetoencephalography (MEG), and functional MRI (fMRI) data. The advantage of using information methods is their independence of a specific probabilistic model, thus enabling the quantification of a far wider variety of interactions and events than would be feasible for a parametric model-dependent method. Information theory can recognize both linear and non-linear relationships between variables.

Here, we used features calculated using information theory to substantially improve the classification algorithms of Li response and BD state. By adding the entropies of the calculated electrophysiological features and the mutual information between the features, we improved the prediction accuracy from 81% to 99% when predicting the patient’s response to Li treatment and from 60% to 74% when classifying a BD state. We also classified the LR and NR neurons from the control neurons after Li and VPA treatment using these algorithms. We found that Li treatment makes LR neurons more neurotypical and harder to distinguish from the control neurons. On the other hand, the Li treatment made the NR neurons more distinguishable from the control neurons.

## Materials and Methods

### Previous work

#### BD Cohorts and reprogrammed iPSCs

BD participants were selected as part of an ongoing genetic research^38^ while healthy volunteers or married-in relatives of certain probands served as control subjects. After signing informed consent, all study participants were assessed using a stringent protocol: pairs of experienced clinician-researchers interviewed them. Following the interviews, a second blind panel of expert clinical researchers formed a standardized diagnosis (Research Diagnostic Criteria^39^ and DSM-IV). Lymphocytes obtained from all individuals were immortalized with Epstein-Barr virus (EBV) and reprogrammed to form iPSCs using the Yamanaka episomal vector defined by Okita et al.^40^ These iPSC lines went through validation of quality control management requirements previously described^15^ before further differentiation and characterization into neuronal cells.

#### Neuronal differentiation

The iPSC colonies that met the quality mentioned above were differentiated into primed neural progenitor cells (NPCs) and then into hippocampal DG granule-cell-like neurons as described previously^15,41^. More than 48% of differentiated neurons expressed the Prox1 gene, a proxy for dentate granule cells. These differentiated neurons were infected with the Prox1::eGFP lentiviral vector on day 12 of the post differentiation period. Electrophysiological recordings were performed on the neurons (with a high Prox1::eGFP expression) using the whole-cell patch-clamp technique in 10-45 days post differentiation.

#### Li and VPA treatment

Neuronal cultures were treated chronically with 1 mM LiCl or 1 mM VPA starting 14 days post differentiation until the patch-clamp experiments. These Li or VPA-treated cells were recorded electrophysiologically in between 22-30 post differentiation periods.

#### Electrophysiological recordings

On the 12th day of differentiation, neurons were infected with the Prox1::eGFP lentiviral vector. Neurons were transferred to a recording chamber using a recording medium solution containing (in mM): 10 HEPES, 4 KCl, 2 CaCl2, 1 MgCl2, 139 NaCl, and 10 D-glucose (solution adjusted to 310 mOsm, pH 7.4). Whole-cell patch-clamp recordings were made from DG-like neurons expressing Prox1::eGFP, typically during 22–30 days of differentiation but ranging from 10–45 days. Internal recording solution containing (in mM): 130 K-gluconate, 6 KCl, 4 NaCl, 10 Na-HEPES, 0.2 K-EGTA, 0.3 GTP, 2 Mg-ATP, 0.2 cAMP, 10 D-glucose, 0.15 % biocytin, and 0.06 % rhodamine were used to fill patch electrodes. Internal solution pH and osmolarity were adjusted to physiological values (pH 7.3, 290–300 mOsm) (pipette tip resistance was usually 10-15 MΩ). Signals were enhanced using a Multiclamp700B amplifier (Sunnyvale, California, USA) and recorded using Axon ‘Instruments’ Clampex 10.2 program (Union City, California, USA). The data were collected at a 20 kHz sampling rate and processed using Clampfit-10 and the MATLAB software kit (release 2014b; The MathWorks, Natick, MA, USA). All measurements were made at room temperature.

#### Electrophysiological analysis

Total evoked action potentials: During patch-clamp recordings, in the current-clamp mode, usually, cells were injected with a holding current needed to hold at −60 mV. Current injection steps of 3 pA were given to the patched cells with a duration of 400 ms beginning from ~12 pA below the holding current. Thirty-five steps of current injections were performed. The study discarded the neurons that required more than 50 pA to retain the membrane potential at −60 mV. The total number of action potentials (spikes) in the 35 steps were counted, and this number is referred to as the total evoked action potentials (cell excitability).

##### Sodium/potassium currents

Voltage clamp mode was used to obtain the sodium and potassium currents. Neurons were held at −60 mV, and periodic voltage steps of 400 ms were given between −90 and 80 mV. In general, these currents were normalized by the cell capacitance to compensate for cell size and to represent ion channel density on the membrane. However, in some of the analyses, the non-normalized currents were compared, and these instances are specifically indicated.

##### Sodium currents

The amplitude of the incoming currents in the voltage-clamp mode in different testing potentials were measured. A strong capacitive transient occurs immediately after a depolarization phase, interfering with the measurements (the gating current). The membrane in the voltage clamp can be approximated as a resistor and a capacitor in a parallel electrical setup. The current in the capacitor operates roughly as the derivative of the change in potential (I=C*dVc/dt) during a voltage step and is much stronger than the currents in the resistor during fast transients because the derivate is large in fast changes. As a result, we can assume a capacitive impedance that scales nearly linear with the voltage step for fast transitions in the membrane potential (dVc). We used this as the reference capacitive current by measuring the current with a −10 mV voltage step from −60 mV to −70 mV, where almost no voltage-gated channels are open. We then generalized this uniformly with the voltage step (for example, multiplying the current provided by the −10 mV step by −2 for a 20 mV voltage step) and subtracting this scaled current from the measured almost wholly removed capacitive transient current. We used the sodium current amplitude at specific test potentials such as −20 mV when calculating some of the information features. These specific potentials were chosen according to where significant differences between groups or drug treatment were found and since they represent physiological conditions.

##### Slow and Fast Potassium currents

Potassium currents were divided into fast potassium currents and slow potassium currents. The maximum current immediately after a depolarization step, generally within a few milliseconds, was used to measure the fast potassium current. Slow potassium currents were measured after 400 ms of depolarization. We measured the potassium currents at specific test potentials such as 0 mV and 20 mV when calculating some of the information features. These specific potentials were chosen according to where significant differences between groups or drug treatment were found and also because of their physiological relevance of these specific test potentials.

##### Input conductance

The input conductance was determined by calculating the current with the cell held in voltage-clamp mode at −70 mV and then at −50 mV. The measured input conductance is the difference in currents divided by the change in membrane potential (20 mV).

##### Capacitance

Capacitance was measured during the recordings by Clampex SW.

##### Spike shape features

The first spike was evaluated for calculating the spike shape features (with the lowest injected current needed for eliciting a spike). The spike threshold was the membrane potential that significantly increased the slope of the depolarizing membrane potential leading to a spike (the first maximum in the second derivative of the voltage vs. time). The 5-ms fast AHP and 1-ms fast AHP were calculated as the difference between the threshold for spiking and the value of the membrane potential 5 ms and 1ms, respectively after the potential returned to cross the threshold value at the end of the action potential. The spike height was calculated as the amplitude difference between the maximum membrane potential during a spike and the threshold. Spike width was calculated as the time it took the membrane potential to change from half the spike height in the rising part to half the spike height in the descending part of the spike (full-width at half-maximum). The rise time of a spike was calculated as the time taken by membrane potential to reach the peak value of membrane potential during the spike from the threshold.

#### Information Theory analysis

Neuronal recordings were categorized into seven groups: control, LR, LR with Li treatment, LR with VPA treatment, NR, NR with Li treatment, and NR with VPA treatment. Every electrophysiological measurement (cell capacitance, no. of spikes, sodium and potassium currents) was uniformly binned into ten bins, and the probability distribution over these bins was calculated assuming ergodicity.

Throughout the study, we used two information-related quantities: Entropy and Mutual Information; Entropy can be understood intuitively as a measure of uncertainty in a given variable. The Entropy of a discrete random variable X is defined as follows^42,43^:

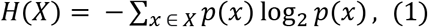

where p(x) is the probability distribution of X. Mutual information can be intuitively understood as a reduction in the uncertainty in one variable by knowing the value of another variable. The mutual information between discrete variables X and Y is defined by^43^:

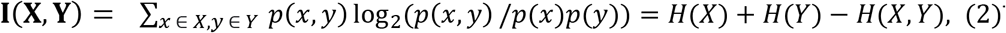

The entropy and mutual information between the electrophysiological features were calculated throughout the neuronal maturation period to follow the developmental trajectory and understand the strength of interaction between the features of the different groups and the changes that occurred by the drug treatment.

##### Correlation and PCA plots

The measured electrophysiological features were normalized by division with the standard deviation. Next, we calculated the covariance matrix between ten physiological features (see Supplementary Table 1) and performed Principle Component Analysis (PCA) using the complete set of features. In the PCA space, we plotted an ellipse for each group that represents the standard deviation of the first PCA component as its width and the standard deviation of the second PCA component as its height. The ellipse’s center was the mean PCA values for the group (“center of mass”).

#### Classification and Prediction

We trained an SVM and random forest classifiers to distinguish between categories based on a set of normalized features derived from the electrophysiological recordings. We performed 10X cross-validation by partitioning the data into ten sections, training the classifier by 90% of the data, and using the remaining data as the test set. Each time, this was done recursively with a different 10% of the data as the test set. Because the entropy depends on the entire dataset distribution, we calculated the entropy individually for each training dataset to avoid cross-talk between the test and training datasets. The Area under the Curve (AUC) in the Receiver Operating Characteristics (ROC) was used to assess the performance of the classification. We calculated the mean accuracy, the mean AUC, and the standard deviation of the separate AUC scores for each test set. Initially, we trained our classifier with a minimal feature set (see Supplementary Table 1). Next, we included six additional spike shape features (see Supplementary Table 1) to train our classifier; lastly, we included 18 additional informational features (see Supplementary Table 1) to further enhance the prediction.

## Results

Only approximately 30% of BD patients were previously shown to respond positively to prophylactic Li treatment^6,7^. We have previously found^15–17^ that BD patients share distinct electrophysiological features, but these divide into subgroups based on the patients’ Li response. Therefore, it is possible to partition the neurophysiology of BD DG granule neurons into two categories according to the Li response of the patient. It is then natural to assume that analyzing the information quantities associated with electrophysiological properties will give further information about the cells and the disorder. The quantities that we chose for this analysis, the entropy and the mutual information, express variation and correlation, respectively, in the dataset.

### Li and VPA differentially affect the entropy of neuronal excitability, depending on the patient’s response to Li

We have previously reported hyperexcitability of BD DG granule neurons^15–18^. We next calculate the entropy of the excitability of the cells. Similar to our previous reports, the analysis performed at 20-30 days post-differentiation shows hyperexcitability of both BD LR and BD NR DG granule neurons compared to controls (Fig. 1a-c presents example traces for control (a), LR (b), and NR (c), and the average excitability in Fig. 1d). The LR neurons become less excitable when treating the neurons with chronic Li treatment (see Methods) (Fig. 1e, neurons 22-30 days’ post differentiation, p=0.02). NR neurons do not change their excitability with Lithium treatment (Fig.1f, p=0.32). VPA treatment decreased the cell excitability both in LR neurons (Fig. 1e, p=0.008) and in NR neurons (Fig. 1f, p=0.04)

**Figure 1.**
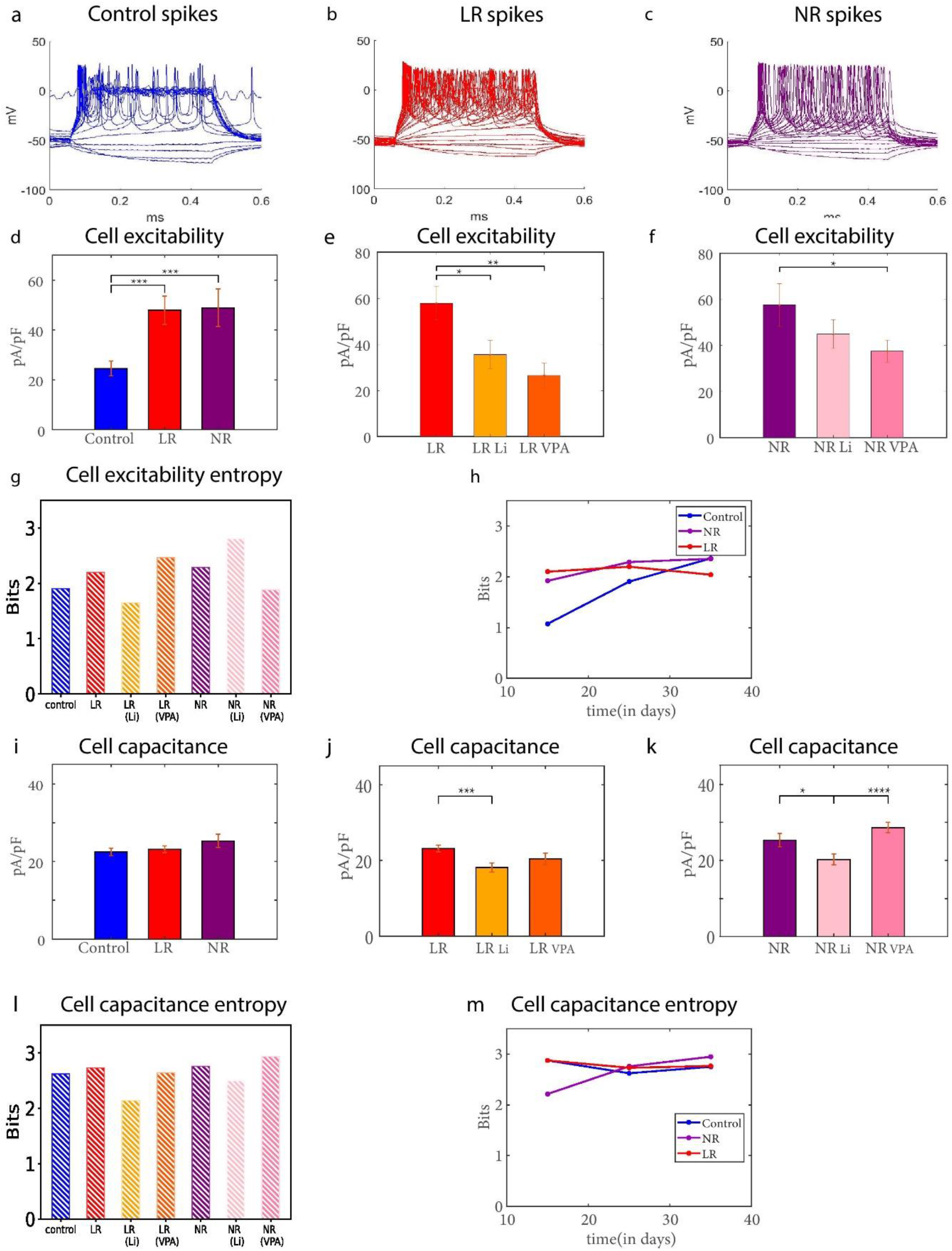
Entropy of Cell excitability and cell capacitance changes differently after Li/VPA treatment in LR and NR DG granule neurons. (The statistics include recordings from control n= 95 neurons, LR n= 84 neurons, LR (Li treatment) n=67 neurons, LR (VPA treatment) n=26 neurons, NR n= 63 neurons, NR (Li treatment) n=47 neurons, NR(VPA treatment) n=75 neurons, all neurons were prox1 positive and therefore are DG granule neurons at 20-30 days post-differentiation unless otherwise stated). (a-c) Example recordings of evoked potentials for control (a), LR (b), and NR(c). (d) Averages of cell excitability for control, LR, and NR DG granule neurons. (e) Averages of cell excitability for LR, LR after Li treatment, and LR after VPA treatment (LR n= 59 neurons, LR (Li treatment) n= 67 neurons, LR (VPA treatment) n=26 neurons for 22-30 days). (f) Averages of cell excitability for NR, NR after Li treatment, and NR after VPA treatment (NR n= 48 neurons, NR (Li treatment) n= 47 neurons, NR (VPA treatment) n=75 neurons for 22-33 days). (g) The entropy of cell excitability for all seven categories (control, LR, LR (Li), LR (VPA), NR, NR(Li), NR(VPA)). (h) The entropy of cell excitability for control, LR, and NR over post differentiation period.(i) Averages of cell capacitance for control, LR, and NR DG granule neurons. (j)) Averages of cell capacitance for LR, LR after Li treatment, and LR after VPA treatment. (k) Averages of cell capacitance for NR, NR after Li treatment, and NR after VPA treatment. (l) The entropy of cell capacitance for all seven categories (control, LR, LR (Li), LR (VPA), NR, NR(Li), NR(VPA)). (m) The entropy of cell capacitance for control, LR, and NR DG granule neurons over post differentiation period. In this figure *p < 0.05, **p < 0.01, ***p < 0.001, ****p < 0.0001 and the error bars show standard errors.

We next examined the differences in the entropy of cell excitability values to assess the diversity between cells within each group. The entropy of cell excitability for BD DG granule neurons was greater than that of the control (2.20 bits for LR, 2.29 bits for NR, and 1.90 bits for the control neurons, Fig. 1g). Notably, the Li treatment decreased the entropy of cell excitability in LR to 1.65 bits (a 25% decrease) while increasing it to 2.81 bits in NR neurons (a 23% increase). With the VPA treatment, the entropy increased (nearly 12%) to 2.46 bits for LR but decreased by 17.8% to 1.88 bits for NR (Fig. 1g).

Interestingly, although Li and VPA reduced the number of spikes in LR and NR neurons, their influence on LR and NR Entropy was in the opposite direction. Furthermore, Li brought the entropy of the neuronal excitability for LR neurons in the direction of the control neurons. At the same time, it had an opposite effect on the NR neurons, increasing the difference of their entropy of excitability from the control neurons.

On the other hand, VPA changed LR entropy of excitability further away from the control neurons. In contrast, it had the opposite effect on the NR neurons, bringing the entropy of the excitability of the NR neurons closer to that of the control neurons. We also calculated the changes that occur in the entropy of excitability during neuronal maturation (Fig. 1h). We noticed that the entropy of cell excitability kept increasing over the maturation period for control neurons but did not change much for the BD neurons suggesting earlier maturation of the BD neurons (Fig. 1h).

The cell capacitance is proportional to the cell surface area and is correlated with neuronal maturation and development. To understand the impact of treatment on the capacitance, we measured the capacitance after Li or VPA treatment. The average cell capacitance was not significantly larger for BD DG granule neurons (p = 0.61 for control vs. LR, p = 0.12 for control vs. NR, Fig. 1i) within 20-30 days (significance is achieved over the entire maturation period of 10-45 days^15^). The Li treatment decreased the capacitance for LR neurons (p=0.00076), and the VPA treatment also had the same trend of reducing the capacitance value for LR (but not statistically significant, p=0.13, Fig.1j). In NR neurons, Li treatment reduced the capacitance (p=0.034); however, VPA did not reduce the capacitance significantly (p = 0.13, Fig. 1k). The entropy of cell capacitance was slightly elevated in NR neurons (2.76 bits) and in LR neurons (2.73 bits) compared to control neurons (2.62 bits). The Li treatment decreased the entropy of cell capacitance to 2.14 bits for LR neurons (down by 21.5%) and to 2.50 bits (down by 9.4%) for NR neurons. VPA treatment did not change the entropy of cell capacitance for LR or NR neurons (a decrease of 3.3% for LR neurons and an increase of 6.4% in NR neurons, Fig. 1l). The entropy of cell capacitance for LR neurons followed the same pattern over post differentiation period as control neurons (Fig. 1m). Still, this entropy started very low in young NR neurons, and, over the maturation period, it grew closer to the entropy of control and LR neurons.

### Li and VPA treatment differentially affects the entropy of the sodium and fast potassium currents depending on the patient’s response to Li

Next, we analyzed the statistics of ionic currents for the three groups (control, LR, and NR) at 20-30 days post-differentiation. Figure 2a-c are example traces of sodium currents at a test potential of −20 mV in control, LR, and NR neurons. Previously, we reported that the sodium current in NR neurons decreased compared to control and LR neurons^15,16^. The average normalized sodium currents at this potential are plotted in Figure 2d, showing a decrease of 36% in NR neurons (p = 0.87 for control vs. LR neurons, p= 0.0087 for control vs. NR neurons, and p= 0.0045 for LR vs. NR neurons). Li treatment increased the sodium currents in the NR DG granule neurons (p=0.04, Fig. 2f), bringing them closer to the controls. VPA treatment almost significantly reduced the sodium currents of the LR neurons (p=0.058, Fig. 2e). The entropy of the sodium current was the highest for the LR neurons (~3 bits). The entropy of the sodium currents in the control neurons was 2.51 bits, and in the NR neurons, it was only 2.26 bits (Fig. 2g). This indicates a narrow distribution of the sodium currents in the NR neurons than control neurons and a broader distribution in the LR neurons.

**Figure 2.**
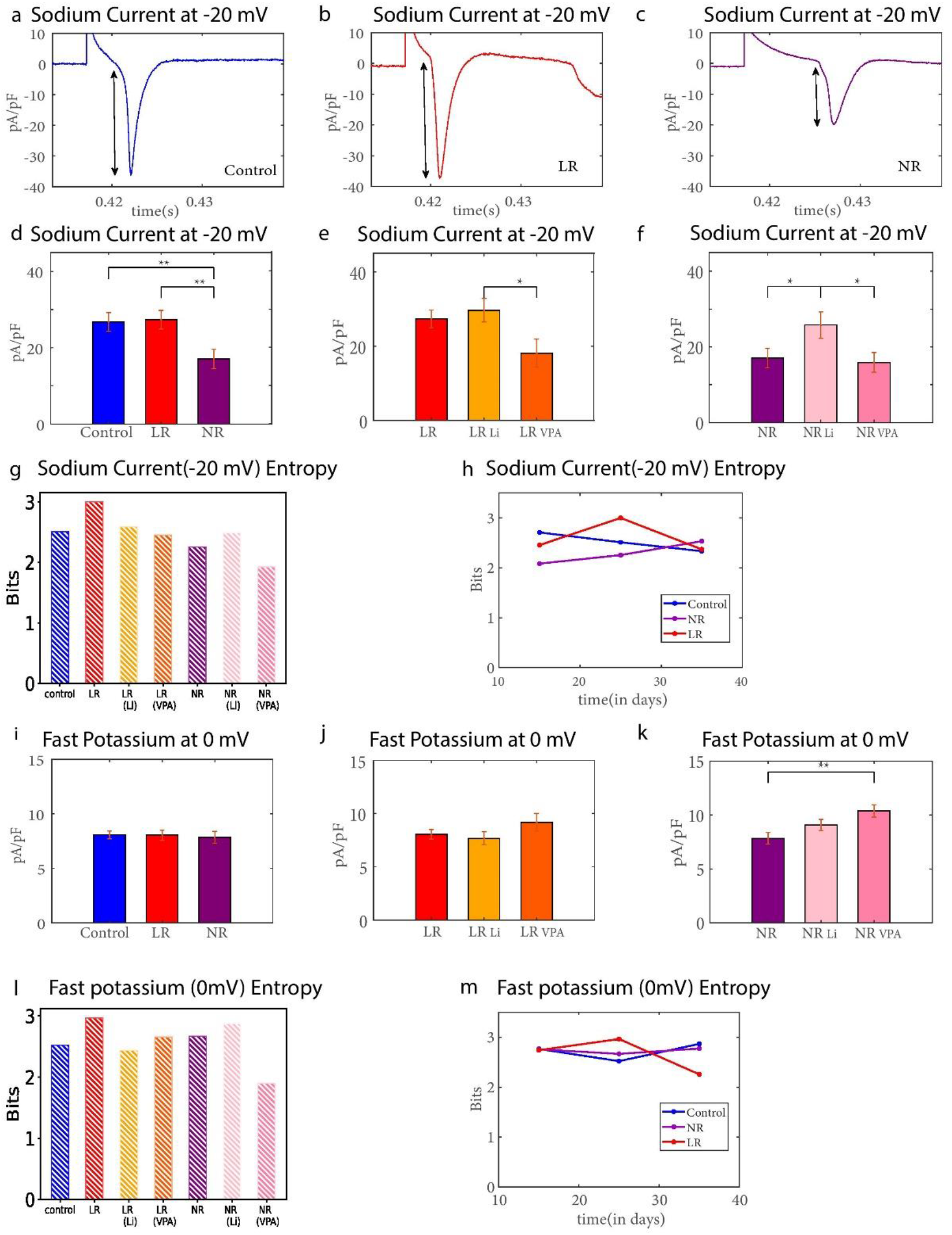
Li treatment showed a contrasting effect on the entropy of sodium and fast potassium currents in LR and NR DG granule neurons. (The statistics include recordings from control n= 95 neurons, LR n= 84 neurons, LR(Li treatment) n=67 neurons, LR (VPA treatment) n=26 neurons, NR n= 63 neurons, NR(Li treatment) n=47 neurons, NR(VPA treatment) n=75 neurons, all neurons were prox1 positive and therefore are DG granule neurons at 20-30 days post-differentiation unless the entire differentiation period is presented).(a-c) Example recordings of Sodium currents at −20 mV for control, LR, and NR. (d) Averages of Sodium currents at −20 mV for control, LR, and NR. (e) Averages of sodium currents at −20 mV for LR, LR after Li treatment, and LR after VPA treatment. (f) Averages of sodium currents at −20 mV for NR, NR after Li treatment, and NR after VPA treatment. (g) The entropy of sodium currents at −20 mV for all seven categories (control, LR, LR (Li), LR (VPA), NR, NR(Li), NR(VPA)). (h) The entropy of sodium currents at −20 mV for control, LR, and NR over the post-differentiation period. (i) Averages of fast potassium currents at 0 mV for control, LR, and NR DG granule neurons. (j) Averages of fast potassium currents at 0 mV for LR, LR after Li treatment, and LR after VPA treatment. (k) Averages of fast potassium currents at 0 mV for NR, NR after Li treatment, and NR after VPA treatment. (l) The entropy of the fast potassium currents at 0 mV for all seven categories (control, LR, LR (Li), LR (VPA), NR, NR(Li), NR(VPA)). (m) The entropy of the fast potassium currents at 0 mV for control, LR, and NR neurons over post differentiation period. In this figure *p < 0.05, **p < 0.01, ***p < 0.001, ****p < 0.0001 and the error bars show standard errors.

The Li treatment reduced the entropy of the sodium current for the LR neurons to 2.59 bits (down by 13.7%, in the direction towards the control neurons, Fig. 2g) while increasing the NR entropy of the sodium current to 2.49 bits (up by 9.5%, in the direction towards the control neurons). After VPA treatment, this entropy decreased to 2.45 bits for LR neurons (~ 18%), and it also decreased to 1.93 bits for NR neurons (14.6%) (Fig. 2g). The entropy of the sodium currents monotonically decreased throughout the differentiation for control neurons (2.7084 for 10-20 days, 2.5113 for 20-30 days, and 2.3338 for 25-45 days, Fig. 2h) while it monotonically increased for NR neurons (2.0839 for 10-20 days, 2.2550 for 20-30 days and 2.5331 for 25-45 days). However, in the LR neurons, the entropy of the sodium current initially increased and then decreased (Fig. 2h).

We next calculated the fast potassium currents at different test membrane potentials. Figure 2i shows the mean values of the fast potassium current at a test potential of 0 mV in control, LR, and NR neurons. The fast potassium current did not change between control and BD neurons (p = 0.96 for control vs. LR neurons, p = 0.72 for control vs. NR neurons). The Li treatment did not cause a significant increase in the fast potassium currents of NR neurons (p=0.11). In comparison, VPA increased the fast potassium currents for NR neurons (for NR neurons p=0.002) but did not cause a significant change for LR neurons (p=0.23). Although the fast potassium current average at 0 mV before treatment was similar between control and BD neurons, the entropy of the fast potassium current was significantly different between control, LR, and NR neurons. The entropy of the fast potassium current was 2.96 bits for LR neurons, compared to 2.67 bits in NR neurons, and compared to 2.52 bits in control neurons. Li treatment decreased the entropy of the fast potassium current in LR neurons to 2.44 bits (a decrease of 17.9%) but increased the entropy of NR sodium current to 2.87 bits (an increase of 7.6%). After VPA treatment, this entropy decreased to 2.66 bits for LR neurons (10.4% reduction) and 1.90 bits for NR neurons (28.9% reduction) (Fig. 2l). The entropy of the fast potassium current over the differentiation period changed similarly for control and NR neurons. However, it exhibited a considerable reduction for LR neurons (Fig. 2m).

### Effects of Li and VPA on the entropy of the slow and fast potassium current

We next analyzed the fast and slow potassium currents at a test potential of 20 mV for neurons at 20-30 days post-differentiation. We chose +20 mV as a test potential in our analysis, as numerous potassium channels open at this potential, yet it is physiological. A representative trace of slow and fast potassium currents at 20 mV is presented in Figure 3a-c for the three groups. The amplitude of the slow potassium current was not significantly different between the three groups (Fig. 3d). The mean value of the slow potassium currents in LR neurons was not significantly different with the chronic application of Li and VPA (Fig. 3e, p=0.2925 for LR vs. LR Li neurons, p = 0.2937 for LR vs. LR VPA neurons). For NR neurons, the slow potassium current also did not significantly change with the chronic application of Li (p=0.2883) but decreased with VPA treatment (Fig. 3f, p= 0.058, close to being significant for NR vs. NR with VPA treatment). While calculating the entropies, we report that NR neurons (2.85 bits) had the highest entropy of the slow potassium current, followed by control neurons (2.64 bits) and LR neurons (2.56 bits)(Fig. 3g). Li treatment increased the entropy of the slow potassium current for LR neurons to 2.85 bits (an increase of 11.3%) and in NR neurons to 2.97 bits (a rise of 4%). After VPA treatment, the entropy increased to 2.91 bits for LR neurons (an increase of 13.9 %) while it decreased to 2.51 bits for NR neurons (a decrease of 12%) (Fig. 3g). The entropy of the slow potassium current for NR and control neurons did not change much over the post-differentiation period, while for LR neurons, it decreased over time (Fig. 3h).

**Figure 3.**
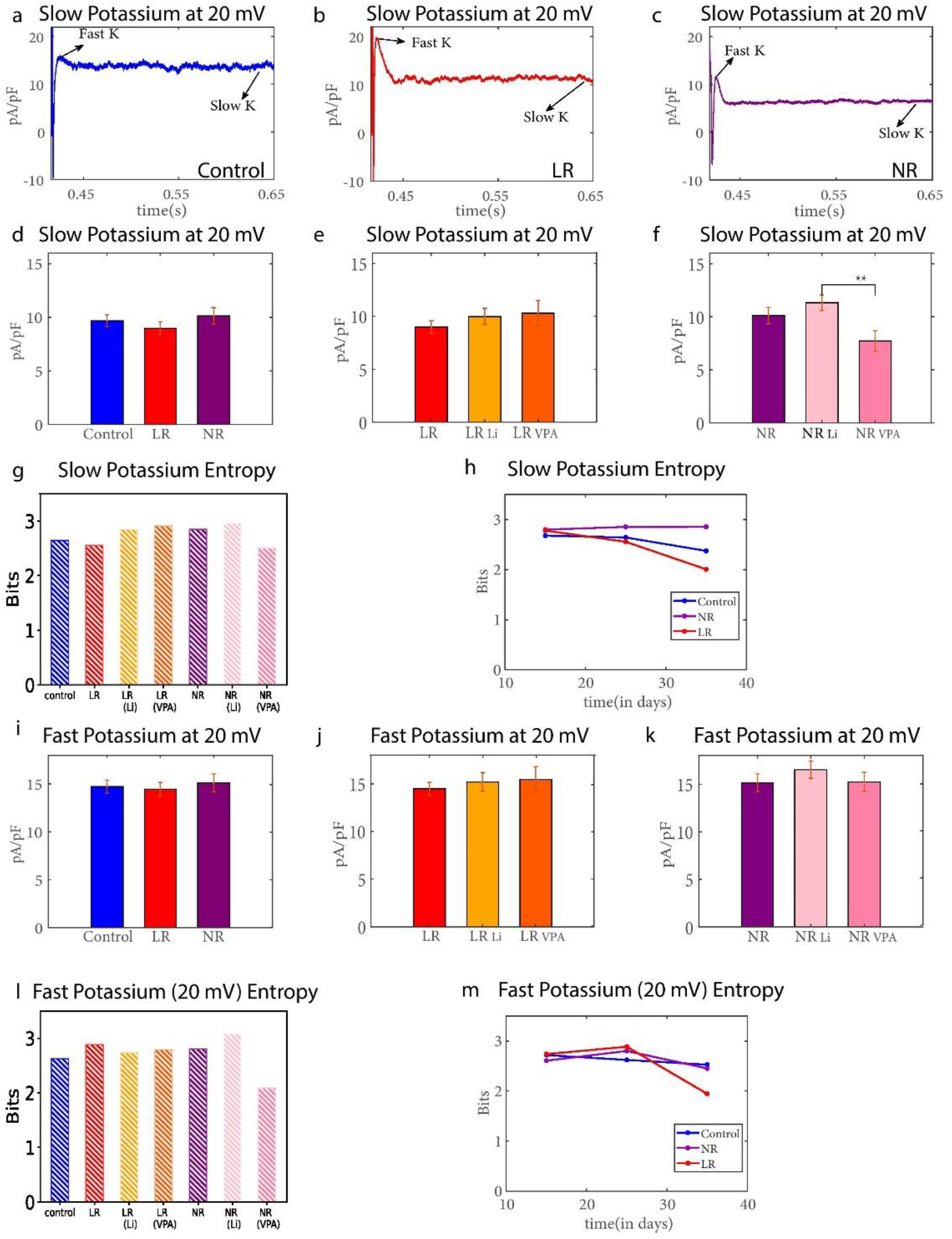
The comparison of entropy of slow and fast potassium currents showed different patterns across LR and NR DG granule neurons after treatment. (The statistics include recordings from control n= 95 neurons, LR n= 84 neurons, LR(Li treatment) n=67 neurons, LR (VPA treatment) n=26 neurons, NR n= 63 neurons, NR(Li treatment) n=47 neurons, NR(VPA treatment) n=75 neurons, all neurons were prox1 positive and therefore are DG granule neurons at 20-30 days post-differentiation unless the entire differentiation period is presented). **(a-c)** Example recordings of slow potassium currents at 20 mV for control, LR, and NR DG granule neurons. **(d)** Averages of slow potassium currents at 20 mV for control, LR, and NR granule neurons. **(e)** Averages of slow potassium currents at 20 mV for LR, LR after Li treatment, and LR after VPA treatment. **(f)** Averages of slow potassium currents at 20 mV for NR, NR after Li treatment, and NR after VPA treatment. **(g)** The entropy of the slow potassium currents at 20 mV for all seven categories (control, LR, LR (Li), LR (VPA), NR, NR(Li), NR(VPA)). **(h)** The entropy of slow potassium currents at 20 mV for control, LR, and NR over the post differentiation period. **(i)** Averages of fast potassium currents at 20 mV for control, LR, and NR. **(j)** Averages of fast potassium currents at 20 mV for LR, LR after Li treatment, and LR after VPA treatment. **(k)** Averages of fast potassium currents at 20 mV for NR, NR after Li treatment, and NR after VPA treatment. **(l)** The entropy of fast potassium currents at 20 mV for all seven categories (control, LR, LR (Li), LR (VPA), NR, NR(Li), NR(VPA)). **(m)** The entropy of fast potassium currents at 20 mV for control, LR, and NR over the post differentiation period. This figure **p < 0.01 and the error bars show standard errors.

We found that the mean value of the fast potassium current at 20 mV was not significantly different across control, LR, and NR neurons. The mean value of this fast potassium current did not change significantly for LR neurons with the application of Li and VPA (p =0.53 for LR vs. LR Li, p = 0.49 for LR vs. LR VPA Fig. 3j). Similarly, it did not change significantly with Li or VPA treatment for NR neurons (p = 0.3 for NR vs. NR Li, p = 0.94 for NR vs. NR VPA Fig. 3k). Calculating the entropy of the fast potassium current across seven categories revealed that LR (2.89 bits) neurons had higher entropy than control neurons (2.62 bits) and NR neurons (2.80 bits), which decreased after chronic Li treatment (2.74 bits) and chronic VPA treatment (2.79 bits) (Fig. 3l). Li treatment for NR neurons increased the entropy of the fast potassium current to 3.08 bits, and interestingly, VPA brought it drastically down to 2.09 bits. The entropy of the fast potassium current in BD DG granule neurons increased initially and then declined, while for the other two groups, it did not change much (Fig. 3m)

### Treatment efficacy can be speculated by analyzing the mutual information properties between electrophysiological features

We characterized the relationships between our seven functional features by mutual information. Mutual information (MI) measures the degree of intrinsically complicated dependencies between the features such as the ionic currents, cell capacitance, and various phenotypes such as hyperexcitability. Looking to see whether treatment will change the features of the neurons to be more neurotypical, we analyzed the mutual information between different combinations of physiological features extracted from our recordings to understand the connections between these values in BD DG neurons. We calculated the mutual information between cell excitability and cell capacitance (MI exc-cap, Fig. 4a(i)), fast potassium current at 0 mV and cell capacitance (MI Fast K (0)-cap, Fig. 4b(i)), fast potassium current at 0 mV and cell excitability (MI Fast K (0)-exc, Fig. 4c(i)), fast potassium current at 0 mV and sodium current (MI Fast K (0)-Na, Fig. 4d(i)), fast potassium current at 20 mV and sodium current (MI Fast K (20)-Na, Fig. 4e(i)), cell excitability and sodium current (MI exc-Na, Fig. 4f(i)), slow potassium current at 20 mV and cell excitability (MI Slow K (20)-exc, Fig. 4g(i)), Sodium current, fast potassium current at 20 mV and cell excitability (MI Na Fast K exc(20), Fig. 4h(i)), Sodium current, slow potassium current at 20 mV and cell excitability (MI Na Slow K exc(20), Fig. 4i(i)). Interestingly, the BD (LR and NR) has a higher MI value for most MI pairs than the controls.

**Figure 4.**
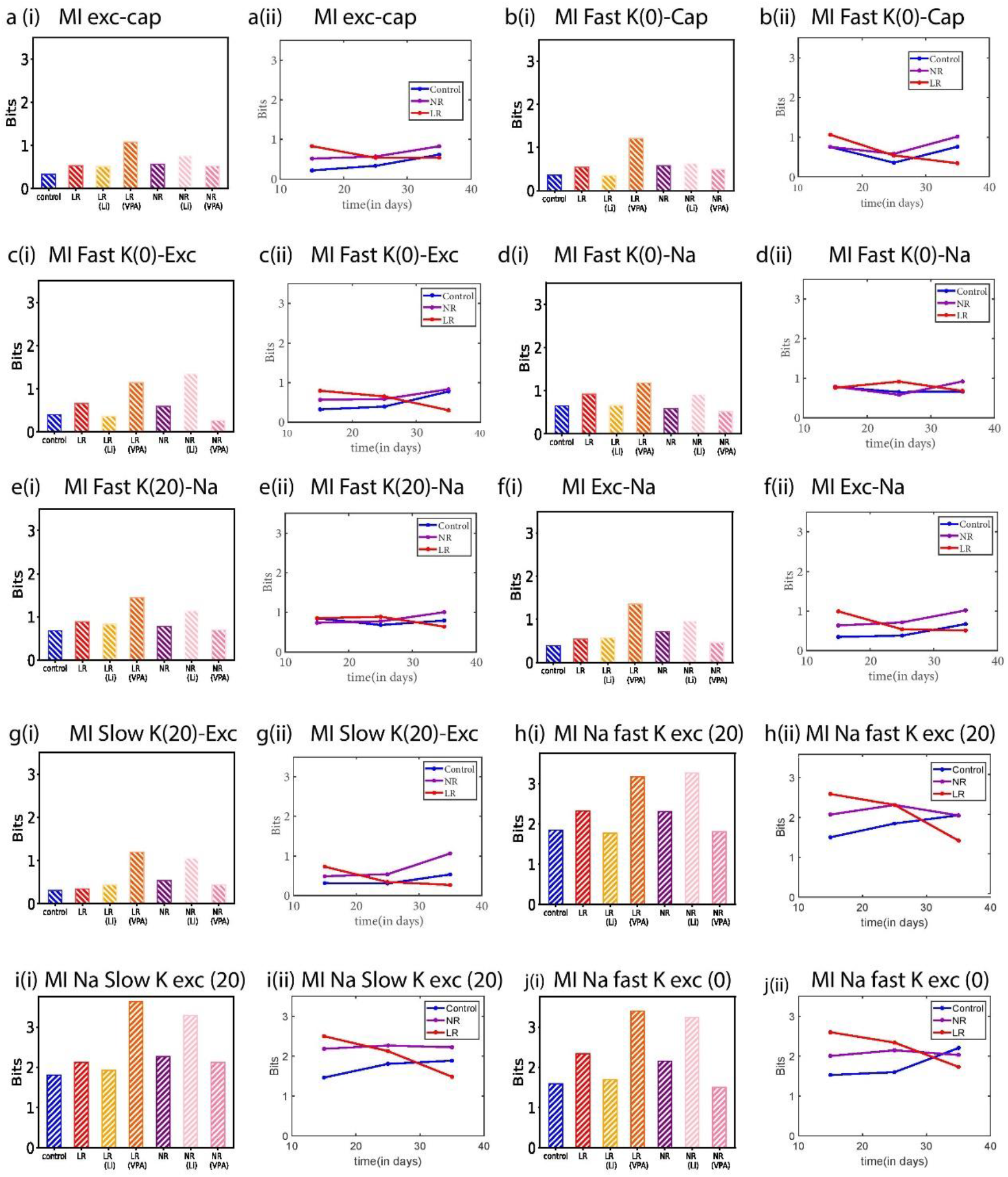
Mutual Information (MI) of different electrophysiological features in all seven categories showed drastic differences between categories (control, LR, NR, with or without Li or VPA treatment). (a) (i) MI of cell excitability and cell capacitance (20-30 days) for all categories and (ii) for control, LR, and NR over the post-differentiation period. (b)(i) MI between fast potassium currents (at 0 mV) and cell capacitance (20-30 days) for all categories and (ii) for control, LR, and NR over the post-differentiation period. (c)(i) MI between fast potassium currents (at 0 mV) and cell excitability (20-30 days) for all categories and (ii) for control, LR, and NR over the post-differentiation period. (d)(i) MI between fast potassium currents (at 0 mV) and sodium currents (at −20 mV) (20-30 days) for all categories and (ii) for control, LR, and NR over the post-differentiation period. (e)(i) MI between fast potassium currents (at 20 mV) and sodium currents (at −20 mV) (20-30 days) for all categories and (ii) for control, LR, and NR over the post-differentiation period. (f)(i) MI between cell excitability and sodium currents (at −20 mV) (20-30 days) for all categories and (ii) for control, LR, and NR over the post-differentiation period. (g)(i) MI between slow potassium currents (at 20 mV) and cell excitability (20-30 days) for all categories and (ii) for control, LR, and NR over the post-differentiation period. (h)(i) MI of sodium currents (at −20 mV), fast potassium currents (at 20 mV), and cell excitability (20-30 days) for all categories and (ii) for control, LR, and NR over the post-differentiation period. (i)(i) MI of sodium currents (at −20 mV), slow potassium currents (at 20 mV), and cell excitability (20-30 days) for all categories, and (ii) for control, LR, and NR over the post-differentiation period.

Additionally, Li treatment in the LR neurons decreased the MI of most pairs of features bringing it closer to the control neurons values, while VPA increased the MI value between most pairs, driving it away from the control neurons and making the neurons less neurotypical. On the other hand, Li treatment in the NR neurons increased the MI value of the pairs mostly, driving it away from the control neurons and making the neurons less neurotypical. In contrast, VPA treatment in NR neurons decreased the MI value of most pairs (refer to supplementary table 2 for percentage difference of individual features value between Control and LR/NR neurons). The evolution of these MI value over the differentiation time can be observed in Figures 4a(ii),b(ii),c(ii),d(ii),e(ii),f(ii),g(ii) & h(ii) respectively.

### The mutual information calculations give further insights than the covariance matrix

Figure 5 presents the covariance matrix between the different features in each group with or without treatment. The Mutual information inherits linear and non-linear relationships between the variables, while the covariance only accounts for the same-direction relationships. Figure 5 (a,b,c) depicts the covariance matrix between different physiological features (cell capacitance, cell excitability, sodium current at −20 mV, fast potassium current at 20 mV, slow potassium current at 20 mV, the spike threshold, the spike height, the rise time, the spike width, and the fast AHP (5ms)) for the three groups (control, LR, and NR). Figures 5(d) and 5(e) present the covariance matrix for LR neurons after Li and VPA treatment, respectively. Similarly, Figures 5(f) and 5(g) present the covariance matrix for NR neurons after Li and VPA treatment, respectively. The covariance matrices were usually in a qualitative agreement with the mutual informational values showing similar relationships between the features of the different groups. For instance, the covariance of the sodium current (as well as for the fast and slow potassium currents) and the cell excitability were very high for LR neurons after VPA treatment which showed a similar increase in the mutual information values after the VPA treatment (Fig. 4c(i), 4f(i), and 4g(i)). In NR neurons, the covariance of the fast and slow potassium current with cell excitability decreased with the use of VPA, similar to mutual information. However, the mutual information performs better than the covariance analysis in depicting how Li treatment brings the values of the MI of the LR group to be closer to the control groups and how VPA further distinguished them from the controls. In the NR, the MI analysis shows better than the covariance analysis how Li draws them further away from the control neurons, while VPA brings them closer.

**Figure 5.**
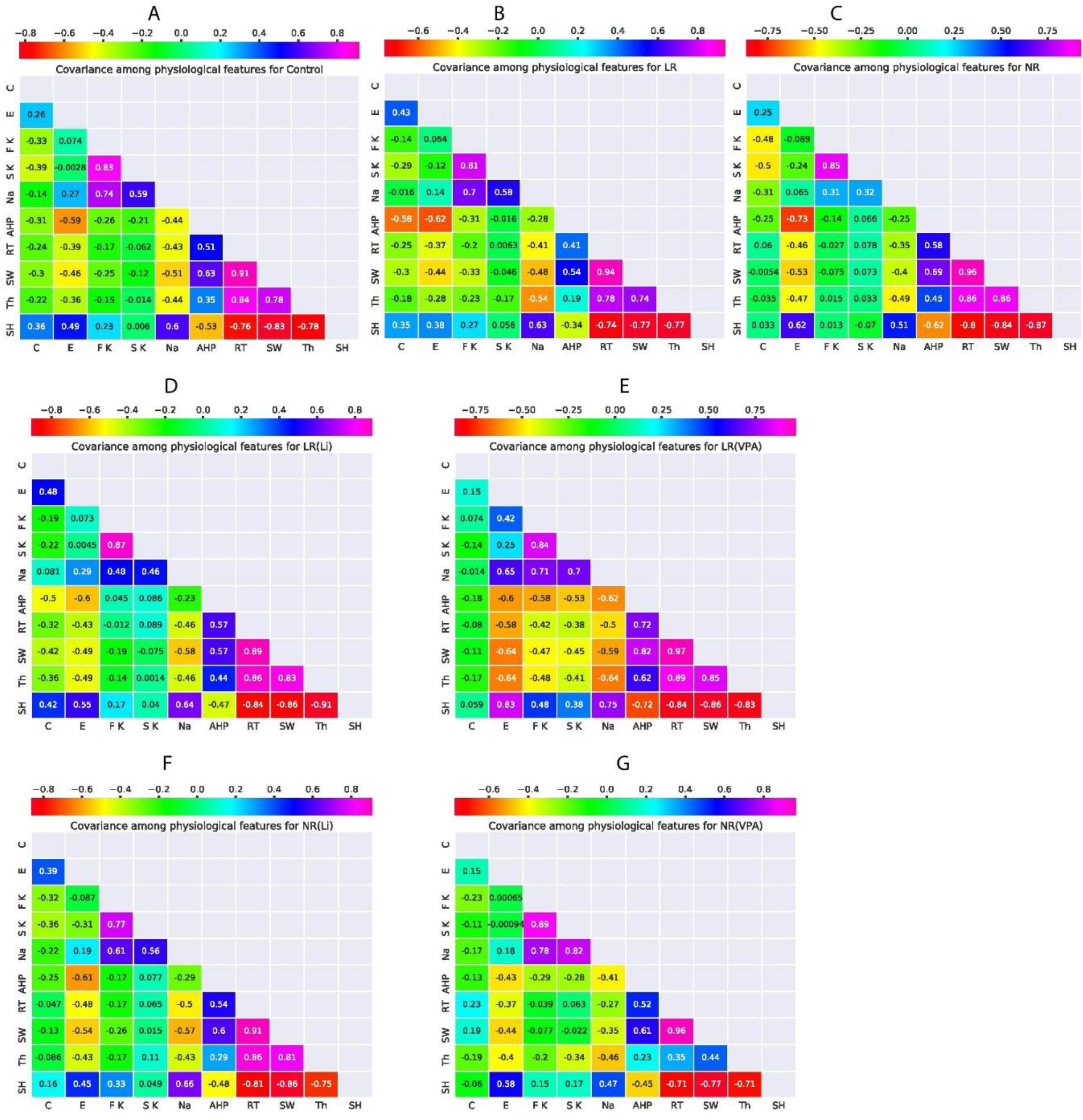
The covariance matrix of ten normalized features extracted from electrophysiological recordings of neurons 20-30 days post-differentiation. In this figure, ‘E’ marks cell excitability, ‘C’ marks cell capacitance, ‘S K’ marks the slow potassium current (at 20 mV), ‘F K’ marks the fast potassium currents (at 20 mV), ‘Na’ marks the sodium currents (at −20 mV), ‘AHP’ marks the fast afterhyperpolarization potential (5 ms), ‘RT’ marks the rise time, ‘SW’ marks the spike width, ‘Th’ marks the spike threshold, and ‘SH’ marks the spike height (amplitude) **(a)** The covariance matrix for control neurons shows that the excitability is highly correlated with the AHP (negatively), spike width (negatively), and spike height. The excitability is also correlated with the sodium current amplitude (p=0.0073). **(b)** The covariance matrix for LR neurons. The correlation between excitability and the fast AHP increased even more compared to control neurons (p=0.21). **(c)** The covariance matrix for NR neurons. The correlation between excitability and the fast AHP increased even more than control and LR neurons. The correlation of excitability with the amplitude of the sodium currents completely diminished (p=0.61). **(d)** The covariance matrix for LR neurons after Li treatment. The correlations of LR excitability with the other features after Li treatment are more similar to the control correlations (sum of squares LR-control is 0.082 and sum of squares LR (Li)-control is 0.072). **(e)** The covariance matrix for LR neurons after VPA treatment. The correlations of LR excitability with the other features after VPA treatment are more similar to the control correlations (sum of squares LR-control is 0.082 and sum of squares LR (VPA)-control is 0.59). **(f)** The covariance matrix for NR neurons after Li treatment. Li treatment made an increase in covariance value between sodium current and excitability(p=0.20). **(G)** The covariance matrix for NR neurons after VPA treatment. The covariance value between both potassium currents and cell excitability decreased after VPA treatment.

### Principal component analysis (PCA) further supports that Li makes LR more neurotypical

In order to assess if drug treatment makes BD neurons more similar to the control neurons, we next performed PCA, including the information features for LR and NR neurons after Li or VPA treatment. Figure 6 presents the analysis with the center of mass of the PCA values for each LR and NR neuron. These are plotted before and after Li/VPA treatment for LR and NR neurons as well as for control neurons. The ellipse around the center of mass represents the standard deviation of the PCA eigenvectors. Figures 6a and 6b present the control, LR, and LR after treatment (Li or VPA, respectively).

**Figure 6.**
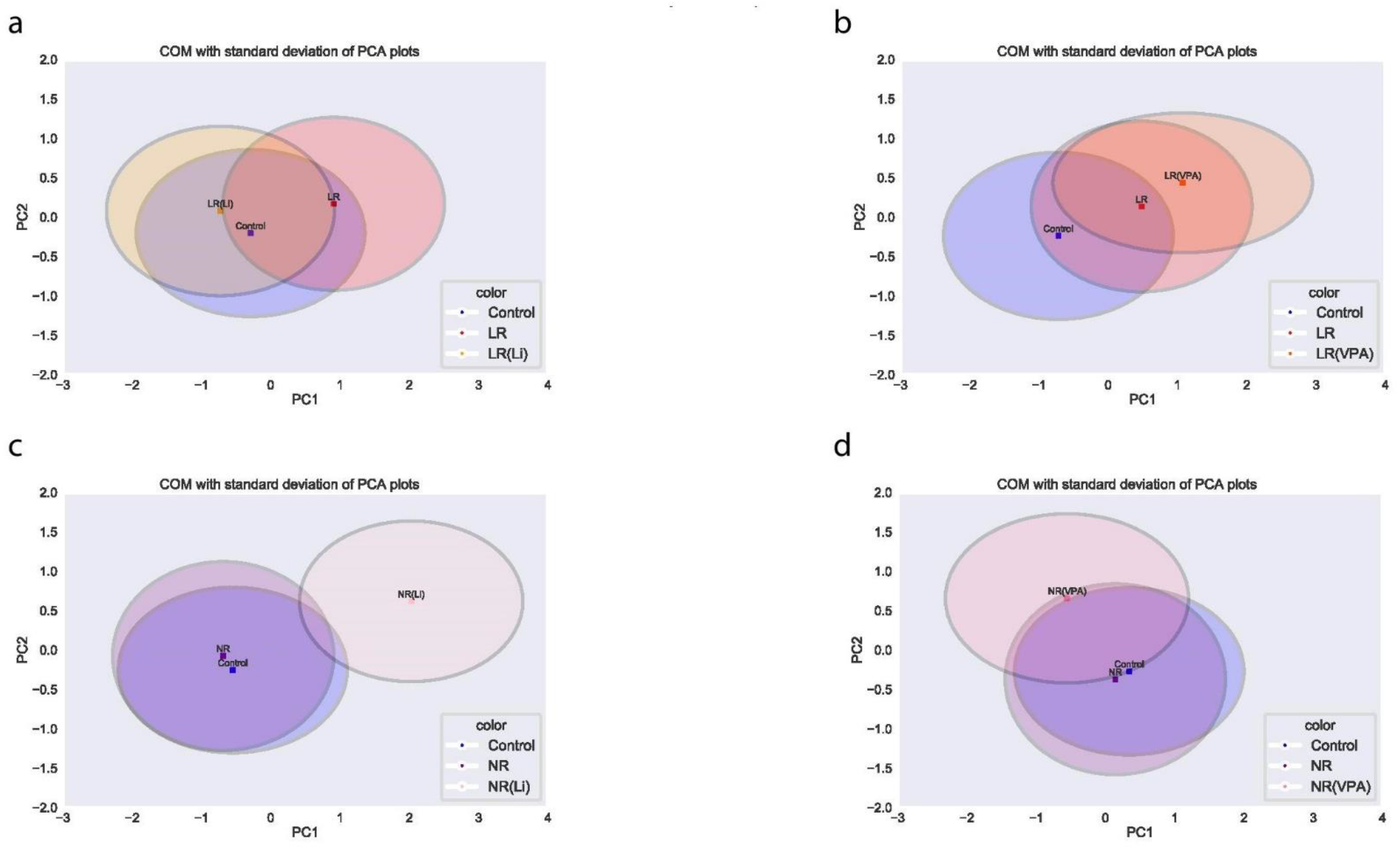
Mean PCA values with standard deviation reflect treatment efficacy in BD neurons. Ellipses centered at mean PC1 and PC2 values and constructed from the standard deviation of PC1 and PC2 values using a set of all features for three groups: control (Cl), LR, and **(a)** LR (after Li treatment)**(b)** LR (after VPA treatment. Again ellipses centered at mean PC1 and PC2 values and constructed from the standard deviation of PC1 and PC2 values using a set of all features for three groups: control (Cl), NR, and **(c)** NR (after Li treatment) **(d)** NR (after VPA treatment).

Similarly, figures 6c and 6d display PCA values for the control, NR, and NR after treatment (Li and VPA, respectively). These plots show that in the PC1/PC2 space, Li treatment has the effect of changing LR neurons to be more similar to control neurons, while VPA moves them further away in this space. Both Li and VPA drive NR neurons further away from the control neurons in the PC1/PC2 space.

### Adding Information theory features improves the classification of BD state substantially

We next checked whether the additional information features would help us classify better control vs. BD and Li response of BD patients using a random forest or a support vector machine (SVM) classifiers (see Material and Methods for more details). When using a random forest classifier with a 10X cross-validation for classification, the mean accuracy when predicting BD state (vs. control) was 62%, with a poor Area Under the Curve (AUC) score (mean AUC (mAUC) = 0.63 ± 0.15) using the Receiver Operating Characteristic (ROC) (Fig. 7a). After including spike shape features, the mean accuracy was 60%, with a poor AUC score (mAUC = 0.63 ± 0.17) (Fig. 7b). When additionally including the information theory features (refer to Material and Methods for more details), the mean accuracy increased to 74% with a good AUC score (mAUC = 0.84 ± 0.05) (Fig. 7c).

**Figure 7.**
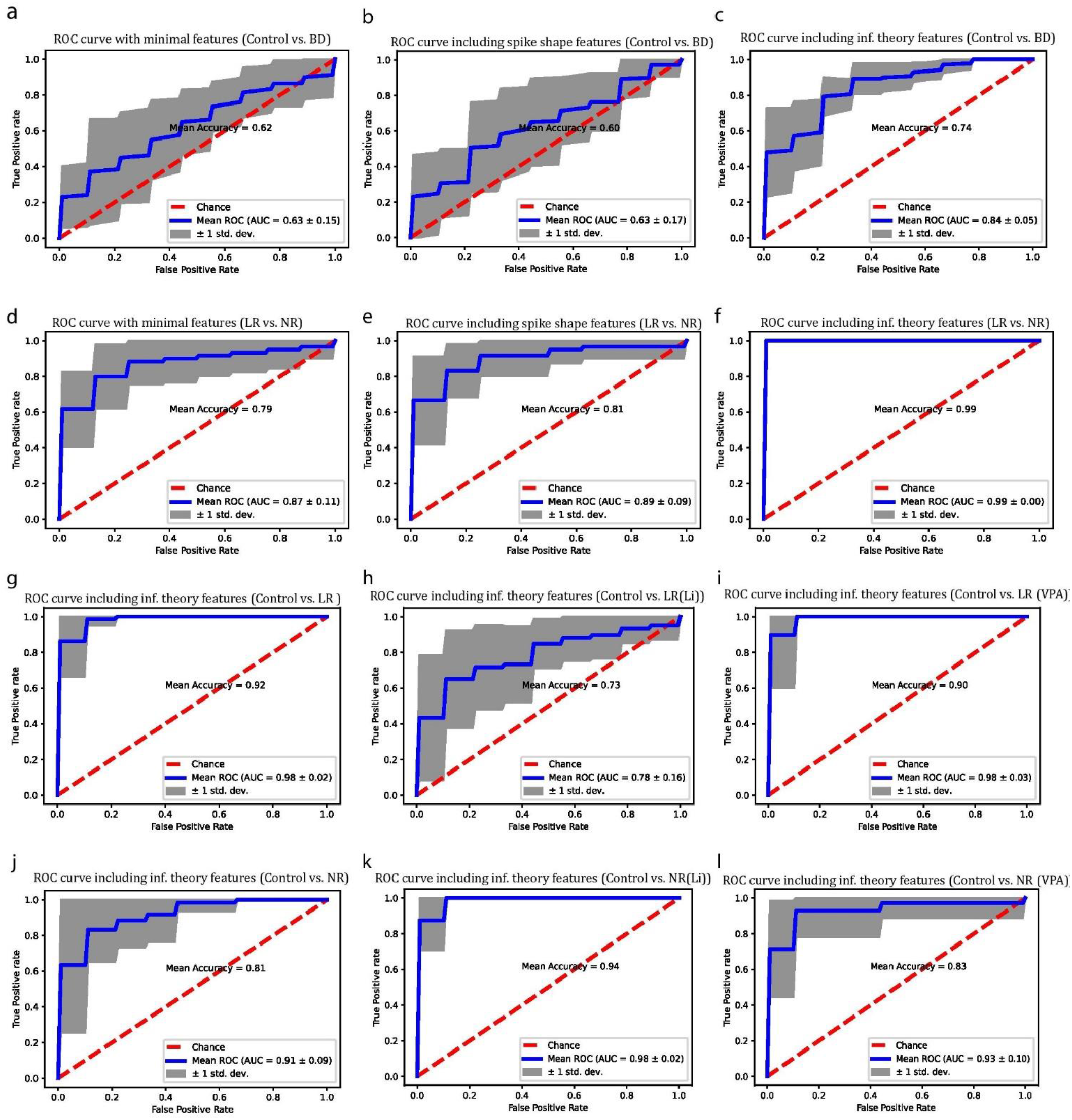
Classifier trained, including mutual information features, showed the highest precision and accuracy. ROC plot when classifying between control and BD neurons after training a random forest classifier with **(a)** Minimal features set **(b)** Including spike shape features **(c)** Including the entropy and mutual information (see Supplementary Table 1). ROC plot between LR and NR neurons after training a random forest classifier with **(d)** Minimal features set **(e)** including spike shape features **(f)** Including the entropy and mutual information. Next, we used an SVM classifier (with the entire features set including the entropy and mutual information) to determine treatment efficacy and plotted the ROC between **(g)** control and LR neurons **(h)** control and LR neurons after Li treatment **(i)** control and LR neurons after VPA treatment. Similarly, we plotted the ROC between **(j)** control and NR neurons **(k)** control and NR neurons after Li treatment **(l)** control and NR neurons after VPA treatment.

### Adding Information theory features further leads to better prediction of Li treatment efficacy in the BD patients

Figures 7d, 7e, and 7f display the prediction between LR vs. NR neurons results in three training options: with the minimal feature set, including spike shape features, and the complete set, which also includes all the information theory features, respectively. When using the minimal feature set, the mean prediction accuracy between LR vs. NR neurons was 79% (mAUC=0.87 ± 0.11). When adding spike shape features, the accuracy increased to 81% (mAUC=0.89 ± 0.09), and when adding the information theory features, the accuracy was 99% (mAUC=0.99 ± 0.00).

To understand how treatment affects the neurons, we next used SVM classification (with the complete feature set including information theory features) to distinguish the patients from the control individuals before and after Li or VPA treatment. Figure 7g shows the classification between control and LR neurons, and Figures 7h and 7i show the classification between control and LR neurons after Li and VPA treatment, respectively. The mean accuracy decreased from 92% (mAUC = 0.98 ± 0.02) (LR vs. control) to 73% (mAUC = 0.78 ± 0.16) after the neurons were treated chronically with Li, suggesting that Li treatment makes the neurons more similar to the control neurons. After treating the LR neurons chronically with VPA, the neurons were still very distinguishable from the control neurons (mean accuracy of 90% with mAUC=0.98 ± 0.03), suggesting that VPA did not make the LR neurons more neurotypical.

Figures 7j, 7k, and 7l show classification between control and NR neurons before and after treatment with Li or VPA. Li treatment increased the classifying accuracy between control and NR neurons from 81% (with mAUC = 0.91 ± 0.09) to 94% (with mAUC = 0.98 ± 0.02), suggesting that the Li makes NR neurons less neurotypical, while VPA also did not change the classification accuracy (mean accuracy of 83% with mAUC = 0.93 ± 0.10), suggesting that it also does not make them more neurotypical.

## Discussion

Identifying the responsiveness of BD patients to Li and VPA at an early stage is critical for providing good treatment to the patients. Furthermore, while drugs are often regarded as effective or not effective by their ability to diminish mood episodes, this may not be the optimal way to assess them, and we would have liked to have a way to assess if the treatment brings them closer to being neurotypical. It was previously reported^15^ that a prediction of BD patients’ response to Li treatment based on electrophysiological features is possible with a low error rate. In the study, a Naïve Bayes Classifier (trained by using characteristics derived from electrophysiological measurements for both categories) was proposed. The study developed a prediction algorithm for the response of a new patient (with unknown Li response) to Li treatment with a 92% successful prediction for young neurons. Here we perform analysis using information theory on electrophysiological measurements of patients’ DG neurons, and the addition of information theory-derived features increases the prediction success significantly. This new analysis also enables us to assess how neurotypical the BD patient-derived neurons become after Li or VPA treatment.

We previously have shown that hyperexcitability is an endophenotype of BD neurons^15,16^. Lithium and VPA reduce excitability, with Li being specific to LR neurons, and VPA reduces the excitability in both groups. Still, human behavior is complex, and the excitability measure by itself does not fully describe the neurophysiological state of the neuron and is very far from describing behavior. To better understand the complexity of the biophysical aspects of neuronal physiology, there are many more aspects of the physiology that should be measured to understand BD-related mechanisms and the effects of specific treatments such as Li and VPA. The information regarding the distribution of distinct characteristics of neurons, such as the number of spikes or capacitance, is lost while calculating the average of these quantities. As we have previously shown when considering the distribution and dynamic changes in specific parameters, we see, for example, that NR neurons are physiologically unstable^17^. The entropy describes the diversity within the distribution, and we have shown here that it gives us more information about the changes from the neurotypical controls. Furthermore, the interplay of the different cellular features and their impact on each other may also be altered in the patient-derived neurons. We, therefore, included the mutual information between the electrophysiological measures in our new analysis. This new angle of approaching the data now shows us a broader view of the neurophysiological features, allowing us to understand the cellular changes in BD better.

The covariance matrix of the electrophysiological features (see Supplementary Table 1 in the Methods section) showed some agreement in the relationships of the different features with the calculated mutual information values (Fig. 5) but not between all the features. For example, the sodium current and the cell excitability covariance were smaller in NR neurons than in control and LR neurons, but the mutual information of these features was higher in the NR neurons. This point may indicate that some of the features may be strongly affecting each other but do not have a clear linear correlation. However, these relationships may be important to the cell functionality, although they are not measured by a simple correlation relationship. Another example can be observed with the effect of Li and VPA treatment when measuring the relationship between cell excitability and the sodium current. Although both Li and VPA treatment increased the correlation between cell excitability and the sodium current, their effect on the mutual information was opposite. Our analysis shows that the mutual information may better reflect treatment efficacy and thus is more informative than the covariance matrix. Performing PCA (Fig. 6) using entropy and mutual information reveals that Li not only decreases the hyperexcitability in LR neurons but also makes them more neurotypical. It is interesting that VPA reduces the hyperexcitability of both LR and NR neurons but does not make them more neurotypical. This analysis that takes not only the first and second moment (the mean and variance) of the distributions into account, but rather complex interactions that are calculated through information theory, reveals essential connections between electrophysiological features that may be important determinants of neurotypical behaviors of our brain cells.

Our classification abilities between control and BD patients have drastically improved by including the information theory features (Fig. 7a-c). The addition of the information theory features also profoundly improves the prediction of the patient’s response to Li treatment. The information theory features were then added to the classification scheme of the patients’ DG neurons compared to the control neurons with and without Li and VPA treatments. Similar to the PCA analysis, the prediction analysis also showed that Li makes LR DG neurons more similar and less distinguishable from control neurons. At the same time, VPA may reduce their excitability but does not make them less distinguishable from the controls. Both Li and VPA do not make NR neurons more distinguishable than control neurons.

Overall, this new way of analysis may have important implications when deciding on the possible treatment of the patients. Furthermore, this new angle of approach may be a suitable method to analyze the effectiveness of treatment and the overall effect on the patients’ neurons.

## Acknowledgment

This material is based upon work supported by the Zuckerman STEM Leadership Program, ISF grant 1994/21, and ISF grant 3252/21.

## Supplementary results

To check whether the minimal feature set or including spike-shape features helps us display the effectiveness of Li/VPA treatment on LR neurons, we trained the classifier with minimal features and included spike-shape features also (presented in supplementary figure 1a-e). Using only minimal features, classifying accuracy between control and LR dropped down to 75% (with mAUC = 0.85 ± 0.12, Supp. Figure 1a), which did not improve even after including spike-shape features (accuracy =72% with mAUC = 0.83 ± 0.12, Supp. Figure 1a). Classifier trained with minimal features showed similar accuracy in classifying control and LR neurons after administering Li treatment (accuracy =75% with mAUC = 0.81 ± 0.12) which remained almost same even after including spike-shape features (accuracy =75% with mAUC = 0.82 ± 0.10). Classification of control and LR neurons after VPA treatment using the SVM classifier trained with minimal features were higher (accuracy =85%) than without treatment case but AUC was very scattered and low (mAUC = 0.70 ± 0.30). Including spike shape features decreased the accuracy for this classification by 1% (accuracy =84%), but AUC improved by a significant amount (mAUC = 0.83 ± 0.19)

Yet again, to test whether including minimal features or spike-shape features assists us to discern the Li/VPA treatment efficacy in the case of NR neurons, we trained the SVM classifier with minimal features as well as including spike-shape features also (presented in supplementary figure 2a-e). Using only minimal features, classifying accuracy between control and NR dropped down to 73% (with mAUC = 0.73 ± 0.17) which did not improve even after including spike-shape features (accuracy =69% with mAUC = 0.72 ± 0.22). Classifier trained with minimal features showed somewhat less accurate classification(which actually improves after including informational features) in classifying control and NR neurons after Li treatment (accuracy =66% with mAUC = 0.63 ± 0.16) which actually improved after including spike-shape features (accuracy =77% with mAUC = 0.75 ± 0.08). Classification of control and NR neurons after VPA treatment using the SVM classifier trained with minimal features were lower (accuracy =69%) than without treatment case but AUC was a bit improved though more scattered (mAUC = 0.79 ± 0.21). Including spike shape features increased the accuracy for this classification by 7% (accuracy =76%) along with improved AUC (mAUC = 0.86 ± 0.12).

**Supplementary Table 1:** Features used in different algorithm.

Features used in different algorithm
| Available Features | Minimal features set | Additional spike shape features | Additional informational features | Covariance plots |
| --- | --- | --- | --- | --- |
| cell capacitance | ✓ | ✓ | ✓ | ✓ |
| cell excitability | ✓ | ✓ | ✓ | ✓ |
| Na current | ✓ | ✓ | ✓ | ✓ |
| Fast K current(0 mV) | ✓ | ✓ | ✓ |  |
| Fast K current(20 mV) | ✓ | ✓ | ✓ | ✓ |
| Slow K current (20 mV) | ✓ | ✓ | ✓ | ✓ |
| Spike Height |  | ✓ | ✓ | ✓ |
| Spike Width |  | ✓ | ✓ | ✓ |
| Spike threshold |  | ✓ | ✓ | ✓ |
| Rise time |  | ✓ | ✓ | ✓ |
| Fast AHP(5 ms) |  | ✓ | ✓ | ✓ |
| Fast AHP (1 ms) |  | ✓ | ✓ |  |
| E cap |  |  | ✓ |  |
| E fast K (20 mV) |  |  | ✓ |  |
| E fast K (0 mV) |  |  | ✓ |  |
| E slow K (20 mV) |  |  | ✓ |  |
| E Na (-20 mV) |  |  | ✓ |  |
| E spikes |  |  | ✓ |  |
| MI Na cap |  |  | ✓ |  |
| MI exc cap |  |  | ✓ |  |
| MI fast K cap (0) |  |  | ✓ |  |
| MI fast K Na (0) |  |  | ✓ |  |
| MI fast K exc (0) |  |  | ✓ |  |
| MI fast K cap (20) |  |  | ✓ |  |
| MI fast K Na (20) |  |  | ✓ |  |
| MI fast K exc (20) |  |  | ✓ |  |
| MI Na exc |  |  | ✓ |  |
| MI Na fast K(20 mV) spikes |  |  | ✓ |  |
| MI Na fast K(0 mV) spikes |  |  | ✓ |  |
| MI Na slow K spikes |  |  | ✓ |  |

**Supplementary figure 1.**
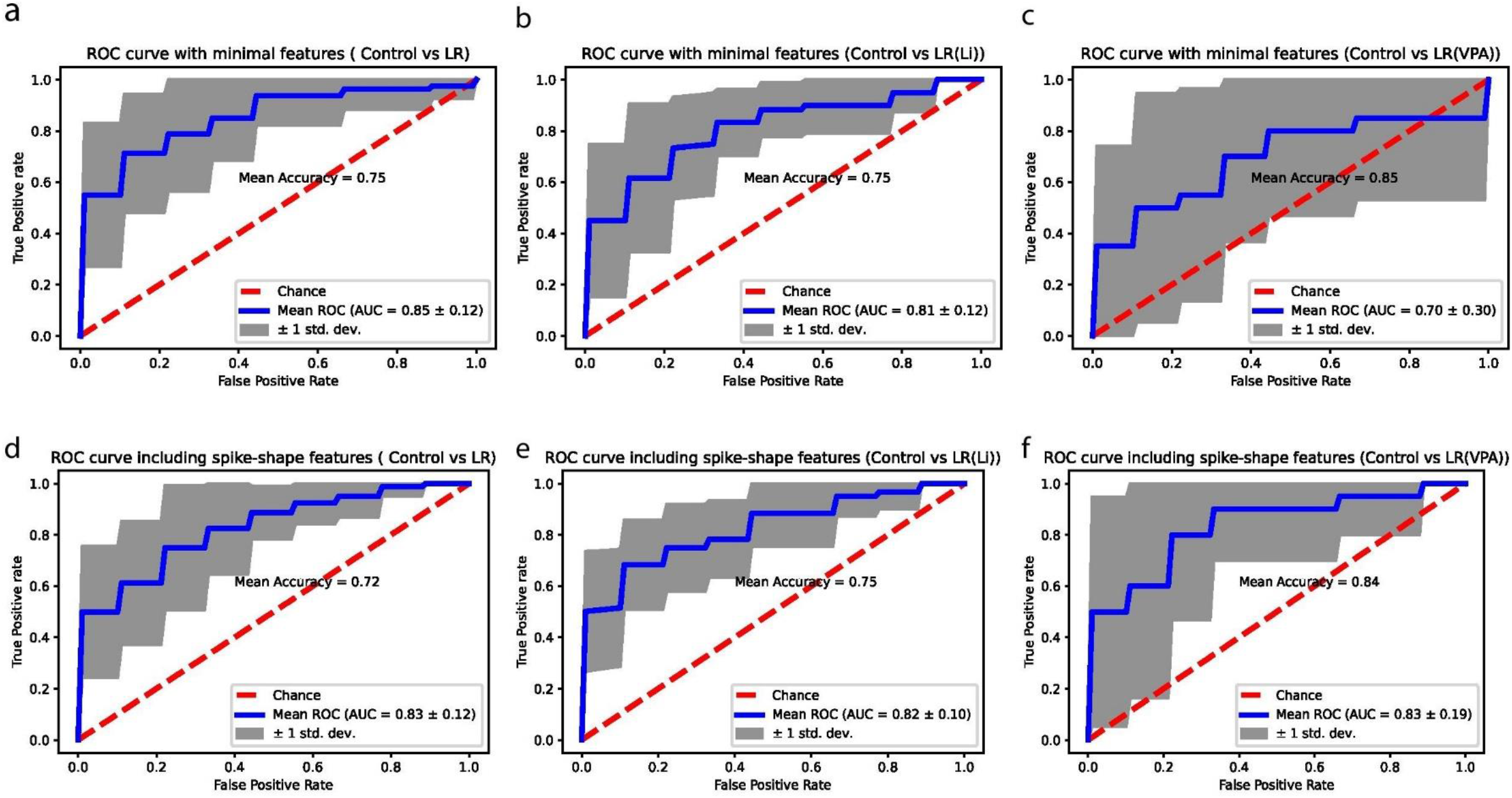
Classification based on minimal features and spike shape features does not reflect treatment efficacy in LR neurons. ROC plot after training an SVM classifier with minimal features set when classifying between **(a)** control and LR (b) control and LR (after Li treatment) **(c)** control and LR (after VPA treatment). ROC plot after training an SVM classifier including spike-shape features when classifying between **(a)** control and LR (b) control and LR (after Li treatment) **(c)** control and LR (after VPA treatment).

**Supplementary figure 2.**
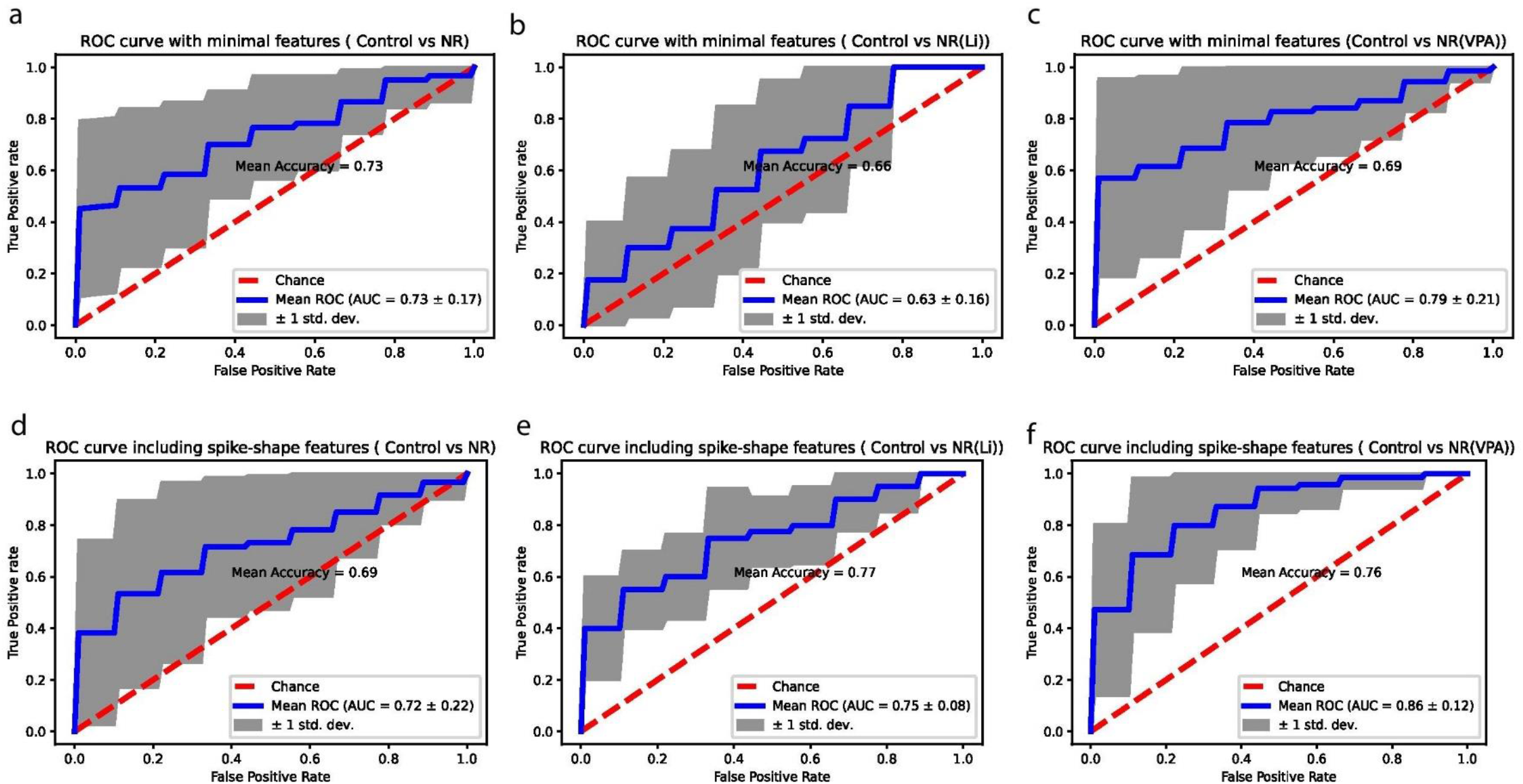
Classification based on minimal features and spike shape features does not reflect treatment efficacy in NR neurons. ROC plot after training an SVM classifier with minimal features set when classifying between **(a)** control and NR (b) control and NR (after Li treatment) **(c)** control and NR (after VPA treatment). ROC plot after training an SVM classifier, including spike-shape features when classifying between **(a)** control and NR (b) control and NR (after Li treatment) **(c)** control and NR (after VPA treatment).

**Supplementary Table 2.** Percentage difference in BD neurons from control before and after treatment.

| Measures<br>(for recordings of 20-30 days Post differentiation period) | Differences in LR(84 neurons) and control(95 neurons) | Differences in LR (After Li) (67 neurons) and control(95 neurons) | Differences in LR (After VPA) (26 neurons) and control(95 neurons) | Differences in NR(63 neurons) and control(95 neurons) | Differences in NR (After Li) (47 neurons) and control(95 neurons) | Differences between NR (After VPA) (75 neurons) and control(95 neurons) |
| --- | --- | --- | --- | --- | --- | --- |
| E cap | 4.10% | -18.30% | 0.69% | 5.18% | -4.70% | 11.93% |
| E fast K (20 mV) | 10.08% | 4.50% | 6.28% | 6.81% | 17.62% | -20.23% |
| E fast K (0 mV) | 17.56% | -3.44% | 5.31% | 5.76% | 13.85% | -24.77% |
| E slow K (20 mV) | -3.26% | 7.66% | 10.14% | 7.96% | 12.30% | -5.03% |
| E Na (-20 mV) | 19.42% | 3.06% | -2.30% | -10.21% | -0.76% | -23.31% |
| E spikes | 15.40% | -13.51% | 29.28% | 20.17% | 47.60% | -1.24% |
| MI Na cap | 125.96% | 78.34% | 223.78% | 77.86% | 81.80% | 52.99% |
| MI exc cap | 61.13% | 54.67% | 225.15% | 70.63% | 129.43% | 56.91% |
| MI fast K cap (0) | 51.79% | -2.91% | 239.57% | 62.58% | 77.24% | 38.65% |
| MI fast K Na (0) | 40.78% | 0.34% | 80.83% | -10.78% | 40.41% | -19.77% |
| MI fast K exc (0) | 65.27% | -8.66% | 188.86% | 48.56% | 240.61% | -32.76% |
| MI fast K cap (20) | 5.66% | 19.06% | 105.36% | 62.83% | 45.08% | 16.50% |
| MI fast K Na (20) | 30.79% | 21.90% | 112.14% | 13.59% | 67.77% | 1.85% |
| MI fast K exc (20) | 55.27% | 24.22% | 220.21% | 84.88% | 200.00% | -5.96% |
| MI slow K cap | -14.96% | 16.91% | 174.98% | 91.70% | 84.30% | 64.63% |
| MI slow K Na | 14.24% | 25.43% | 109.95% | 14.85% | 76.36% | 52.55% |
| MI slow K exc | 12.62% | 41.34% | 291.57% | 76.43% | 246.62% | 41.28% |
| MI Na exc | 41.87% | 48.24% | 256.22% | 87.22% | 151.89% | 19.78% |
| 654<br>MI Na fast<br>K(20 mV)<br>spikes | 25.32% | -4.04% | 71.65% | 25.23% | 77.06% | -2.12% |
| MI Na fast<br>K(0 mV)<br>spikes | 46.63% | 5.57% | 112.48% | 34.52% | 102.65% | -5.74% |
| MI Na slow K<br>spikes | 17.71% | 6.67% | 101.23% | 25.66% | 82.38% | 17.41% |

